# A novel CRISPR-based malaria diagnostic capable of *Plasmodium* detection, speciation, and drug-resistance genotyping

**DOI:** 10.1101/2020.04.01.017962

**Authors:** CH Cunningham, CM Hennelly, JT Lin, R Ubalee, RM Boyce, EM Mulogo, N Hathaway, KL Thwai, F Phanzu, A Kalonji, K Mwandagalirwa, A Tshefu, SR Meshnick, JJ Juliano, JB Parr

**Author notes:** Corresponding author information: Jonathan B. Parr, MD, MPH; 130 Mason Farm Rd., Chapel Hill, NC 27599; 1-919-843-8132. Co-senior authors (JJ Juliano, JB Parr).

## Abstract

CRISPR-based diagnostics are a new class of highly sensitive and specific assays with multiple applications in infectious disease diagnosis. SHERLOCK, or Specific High-Sensitivity Enzymatic Reporter UnLOCKing, is one such CRISPR-based diagnostic that combines recombinase polymerase pre-amplification, CRISPR-RNA base-pairing, and LwCas13a activity for nucleic acid detection. We developed SHERLOCK assays for malaria capable of detecting all *Plasmodium* species known to cause malaria in humans and species-specific detection of *P. vivax* and *P. falciparum*, the species responsible for the majority of malaria cases worldwide. We validated these assays against parasite genomic DNA and achieved analytical sensitivities ranging from 2.5-18.8 parasites per reaction. We further tested these assays using a diverse panel of 123 clinical samples from the Democratic Republic of the Congo, Uganda, and Thailand and pools of *Anopheles* mosquitoes from Thailand. When compared to real-time PCR, the *P. falciparum* assay achieved 94% sensitivity and 94% specificity in clinical samples. In addition, we developed a SHERLOCK assay capable of detecting the dihydropteroate synthetase (*dhps*) single nucleotide variant A581G associated with *P. falciparum* sulfadoxine-pyrimethamine resistance. Compared to amplicon-based deep sequencing, the *dhps* SHERLOCK assay achieved 73% sensitivity and 100% specificity when applied to a panel of 43 clinical samples, with false-negative calls only at lower parasite densities. These novel SHERLOCK assays have potential to spawn a new generation of molecular diagnostics for malaria and demonstrate the versatility of CRISPR-based diagnostic approaches.

**One-sentence summary:** Novel malaria SHERLOCK assays enabled robust detection, speciation, and genotyping of *Plasmodium spp*. in diverse samples collected in Africa and Asia.

## INTRODUCTION

Newly developed technologies that utilize Clustered, Regularly-Interspaced Palindromic Repeat (CRISPR) systems have the potential to revolutionize infectious disease diagnostic testing.*(1)* SHERLOCK, or Specific High-Sensitivity Enzymatic Reporter UnLOCKing, is a CRISPR-based diagnostic assay that has now been used to detect dengue and Zika viruses with excellent sensitivity and specificity.*(2)* Its simple workflow and robust performance characteristics have enabled multiplexed detection and genotyping of Zika and dengue viruses, with increasingly streamlined protocols that facilitate use at the point-of-care.*(3, 4)* SHERLOCK’s potential is perhaps greatest in low-resource settings where improved, reliable diagnostics are urgently needed for multiple pathogens and for malaria, in particular.

Timely and accurate diagnosis is an important component of malaria control and elimination efforts. The current generation of rapid diagnostic tests (RDTs) that detect *Plasmodium falciparum* histidine-rich protein 2 (PfHRP2) are widely deployed, accounting for 74% of all malaria testing in Africa in 2015, but they have shortcomings.*(5)* Conventional PfHRP2-based RDTs only detect *P. falciparum-* and miss low density infections, under approximately 200 parasites/µL. Because PfHRP2 antigenemia can persist for weeks, they can produce false-positive results well after resolution of infection.*(6)* Additionally, increasing reports of false-negative RDT results due to *P. falciparum* parasites that escape detection due to deletion mutations of the *pfhrp2* and *pfhrp3* genes raise concern that PfHRP2-based RDTs may be threatened in select regions of Africa.*(7, 8)* While microscopy is the traditional gold-standard for malaria diagnosis, it is only sporadically available throughout much of Africa because performance is highly operator dependent, it is labor-intensive, and it requires careful, sustained training of personnel. RDTs that detect alternative antigens such as parasite lactate dehydrogenase (pLDH) are less sensitive and heat-stable. To overcome these challenges, a new generation of field-deployable malaria diagnostics capable of detecting diverse species at low parasite densities is needed.

SHERLOCK combines recombinase polymerase amplification (RPA), in vitro transcription, and RNA target detection using custom-designed CRISPR RNA (crRNA) oligonucleotides and Cas13a derived from the bacteria *Leptotrichia wadei* (LwCas13a). In brief, RPA is performed with primers tagged with a T7 promoter sequence, generating short, double-stranded DNA (dsDNA) amplicons of a target sequence. *In vitro* transcription of the RPA product by T7 polymerase produces single-stranded (ssRNA) targets, which are recognized by base-pairing interactions with LwCas13a:crRNA complex. These CrRNA-ssRNA target base-pairing interactions activate collateral RNAse activity of LwCas13a, which cleaves fluorescent or colorimetric RNA reporter molecules in the reaction and produces a detectable signal. These reaction components can be combined into one or two reactions, with detection on a simple fluorimeter or lateral flow strip.*(3)* The highly specific nature of LwCas13a:crRNA complex formation has now been leveraged to distinguish Zika and dengue virus strains and for human cancer genotyping.

We describe novel malaria SHERLOCK assays that enable robust detection of all five *Plasmodium* species known to be pathogenic to humans and species-specific detection of the primary causes of human disease *P. falciparum* and *Plasmodium vivax*. We apply these assays to laboratory parasite strains, diverse clinical isolates from Africa and Asia, and pools of *Anopheles* mosquitoes collected in the field. We then describe a proof-of-concept assay for detection of a single nucleotide variant (SNV) in the *P. falciparum* dihydropteroate synthase (*dhps*) gene that is associated with resistance to sulfadoxine-pyrimethamine, the primary antimalarial medication used in intermittent preventive treatment of malaria in pregnancy (IPTP) throughout Africa.*(9)*

## RESULTS

### Plasmodium SHERLOCK development overview

We translated SHERLOCK to malaria by designing and screening candidate RPA primers and crRNAs, optimizing assay conditions, and validating them on clinical and mosquito samples collected in diverse sites (**Figure 1**). Assay development included the design and testing of two broad categories of RPA primers and crRNAs. The first design targeted conserved and variable regions of the *Plasmodium* 18S rRNA gene and enabled detection and speciation of malaria in clinical samples and mosquitoes. The second design included engineered crRNA-target mismatches to enable detection of the *P. falciparum* A581G mutation associated with resistance to sulfadoxine-pyrimethamine (**Figure 2**). A range of reaction parameters were tested during pilot testing. While we were unable to replicate the “one-pot” reaction approach (combining RPA, IVT, and Cas13a detection) previously described for viral targets,*(3)* we achieved optimal performance using a two-step reaction format (RPA followed by IVT and Cas13a detection) and 10-50-fold higher crRNA concentrations than those originally described by Gootenberg *et al*.*(2)*

**Figure 1.**
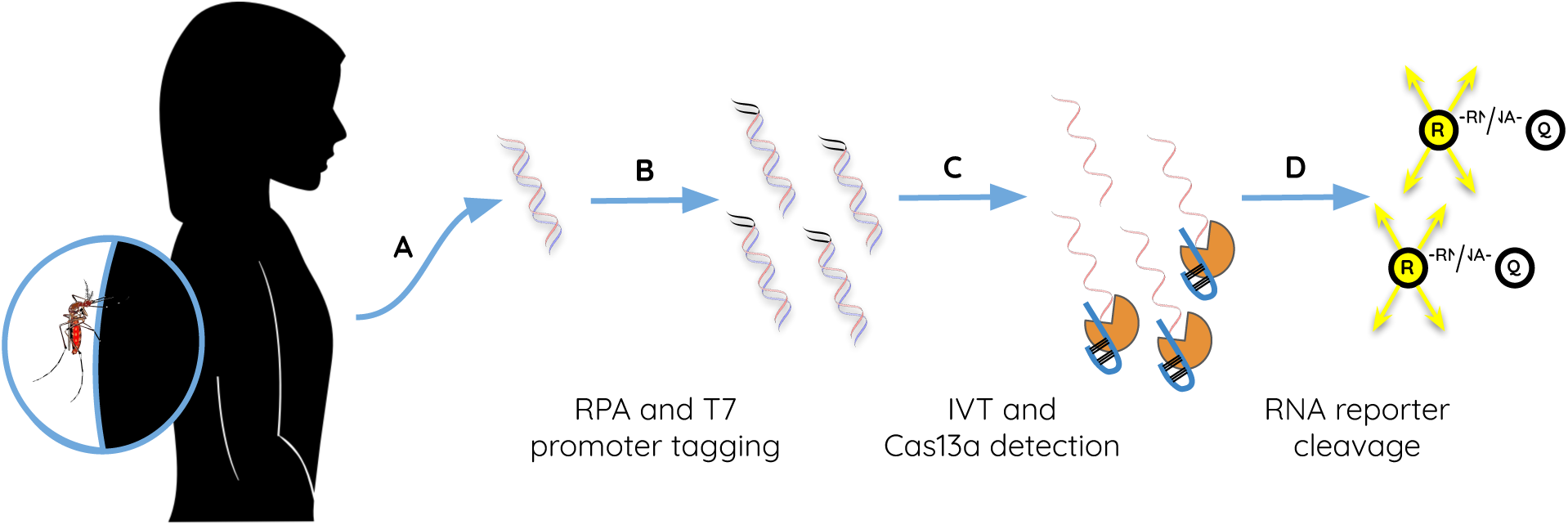
SHERLOCK reaction workflow. (A) DNA extracted from a patient with malaria or infected mosquito is subjected to (B) RPA including T7 promoter-tagged primers for amplification of the *Plasmodium* 18S rRNA or *P. falciparum dhps* genes, followed by (C) IVT and LwCas13a:crRNA complex binding to genus-, species-, or genotype-specific target RNA. This binding triggers collateral (D) activation of LwCas13a RNAse activity and cleavage of reporter RNA, separating the fluorescent reporter from its quencher and producing a signal. Abbreviations: RPA, recombinase polymerase amplification; IVT, in vitro transcription; R, reporter; Q, quencher.

**Figure 2.**
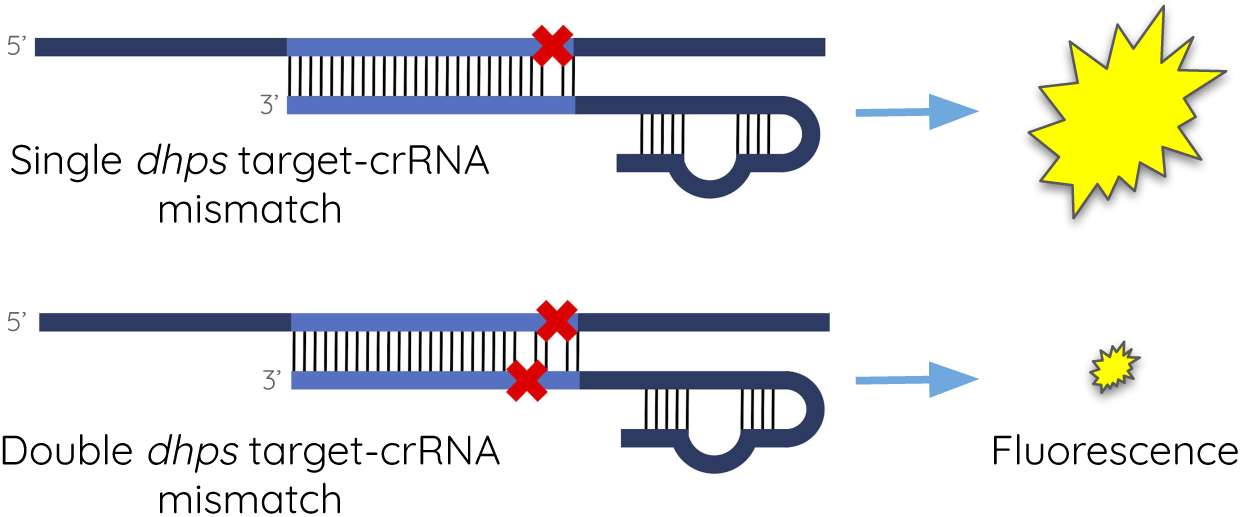
SHERLOCK is capable of distinguishing the *dhps* A581G SNV associated with antimalarial drug resistance. Engineered target-crRNA mismatches produce differential Cas13a activation and fluorescence output.

### Plasmodium SHERLOCK achieves robust analytical sensitivity and specificity

SHERLOCK demonstrated attomolar analytical sensitivity for *Plasmodium* DNA, equivalent to 95% lower limits of detection (LODs) of 2.5-18.8 parasite genomes per reaction (4.2-31.3 aM; **Figure 3, Table 1**) for all three assays. Using only 1 µL of initial DNA input, this is roughly equivalent to 2.5-18.8 parasites/µL. These sensitivities approach those of commonly deployed malaria real-time PCR assays,*(10)* which achieved 0.3-2.5 parasite/µL analytical sensitivities during head-to-head testing in one controlled setting.*(11)* The pan-*Plasmodium* and *P. falciparum* SHERLOCK assays achieved superior LODs compared to *P. vivax*, but all three assays achieved LODs well below those detected by commonly used malaria RDTs (**Table 1**).

**Table 1.** 95% lower limits of detection (LOD) for *Plasmodium spp*. SHERLOCK assays. Input: One μL of DNA template/reaction.

| Assay | LOD (95% CI) | Parasite equivalents/reaction (replicates detected / replicates tested) |  |  |  |  |  |  |  |  |  |
| --- | --- | --- | --- | --- | --- | --- | --- | --- | --- | --- | --- |
|  |  | 10<sup>5</sup> | 10<sup>4</sup> | 10<sup>3</sup> | 10<sup>2</sup> | 20 | 15 | 10 | 5 | 2.5 | 1.25 |
| Pan-*Plasmodium* | 2.5<br>(1.9-21.3) | 3/3 | 3/3 | 3/3 | 3/3 | - | - | 20/20 | 20/20 | 19/20 | 14/20 |
| *P. falciparum* | 6.8<br>(5.3-11.6) | 3/3 | 3/3 | 3/3 | 3/3 | - | - | 20/20 | 17/20 | 9/20 | 8/20 |
| *P. vivax* | 18.8<br>(13.3-146.3) | 3/3 | 3/3 | 3/3 | 3/3 | 20/20 | 20/20 | 10/20 | 19/20 | - | - |

**Figure 3.**
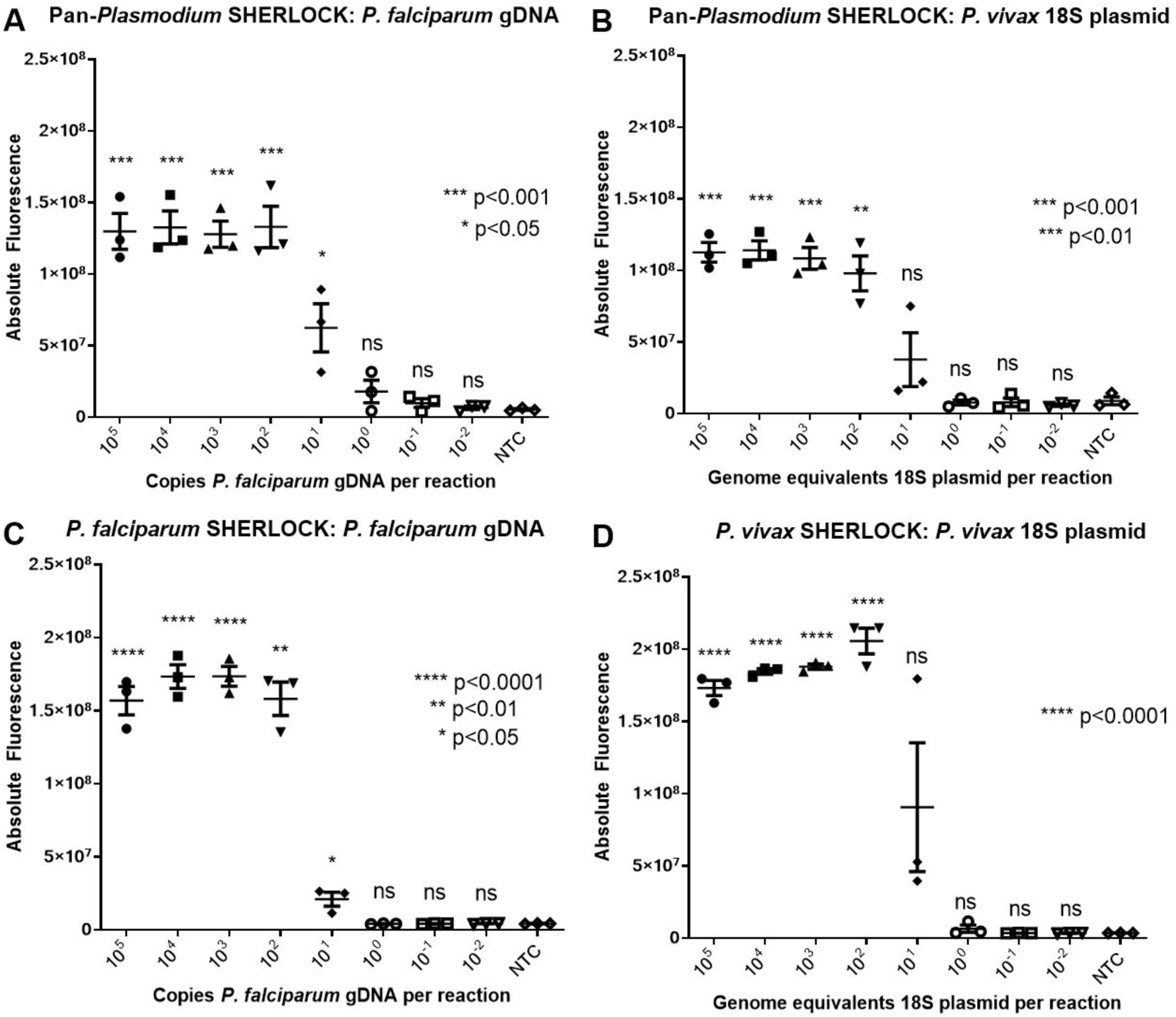
Analytical sensitivity of *Plasmodium spp*. SHERLOCK assays applied to *P. falciparum* strain 3D7 gDNA and *P. vivax* 18S rRNA plasmid DNA. Abbreviations: NTC, no template control.

To test the specificity of the SHERLOCK assays against different *Plasmodium* species, we conducted assays using high concentration 18S rRNA plasmid DNA or gDNA from all five human-infecting *Plasmodium* species (**Figure 4**). The pan-*Plasmodium* SHERLOCK assay detected all five *Plasmodium* species, while the *P. falciparum* assay only displayed activity against *P. falciparum* gDNA. The *P. vivax* SHERLOCK assay detected *P. vivax* 18S rRNA plasmid DNA, but also demonstrated low-level cross-reactivity for *P. knowlesi* gDNA.

**Figure 4.**
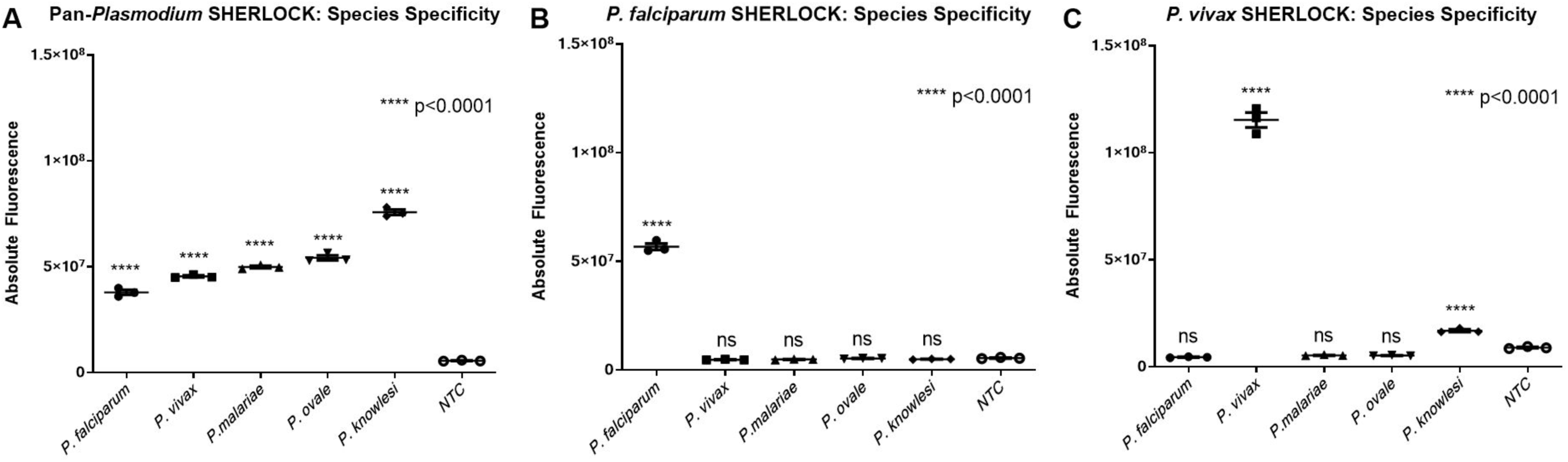
(A) Pan-*Plasmodium*, (B) *P. falciparum*, and (C) *P. vivax* SHERLOCK assays can differentiate *Plasmodium* species. Input for all species was 100,000 genome-equivalents 18S plasmid/reaction or genome-equivalents of gDNA/reaction (*P. knowlesi* only). Abbreviations: NTC, no template control.

### SHERLOCK performs well in clinical samples and infected mosquitoes

SHERLOCK successfully detected and differentiated the four most common human malaria species in clinical samples across a range of parasite densities. When applied to a panel of real-time PCR-positive clinical isolates from the Democratic Republic of the Congo (DRC) and Thailand, the species-specific *P. falciparum* and *P. vivax* assays demonstrated complete clinical specificity (**Figure 5**). When applied to a larger panel of 112 samples from the DRC and Uganda, the *P. falciparum* SHERLOCK assay’s sensitivity and specificity were 94% and 94%, respectively, compared to real-time PCR **(Supplementary Figure 4)**. Threshold Cohen’s kappa was 0.87, consistent with excellent agreement between methods.

**Figure 5.**
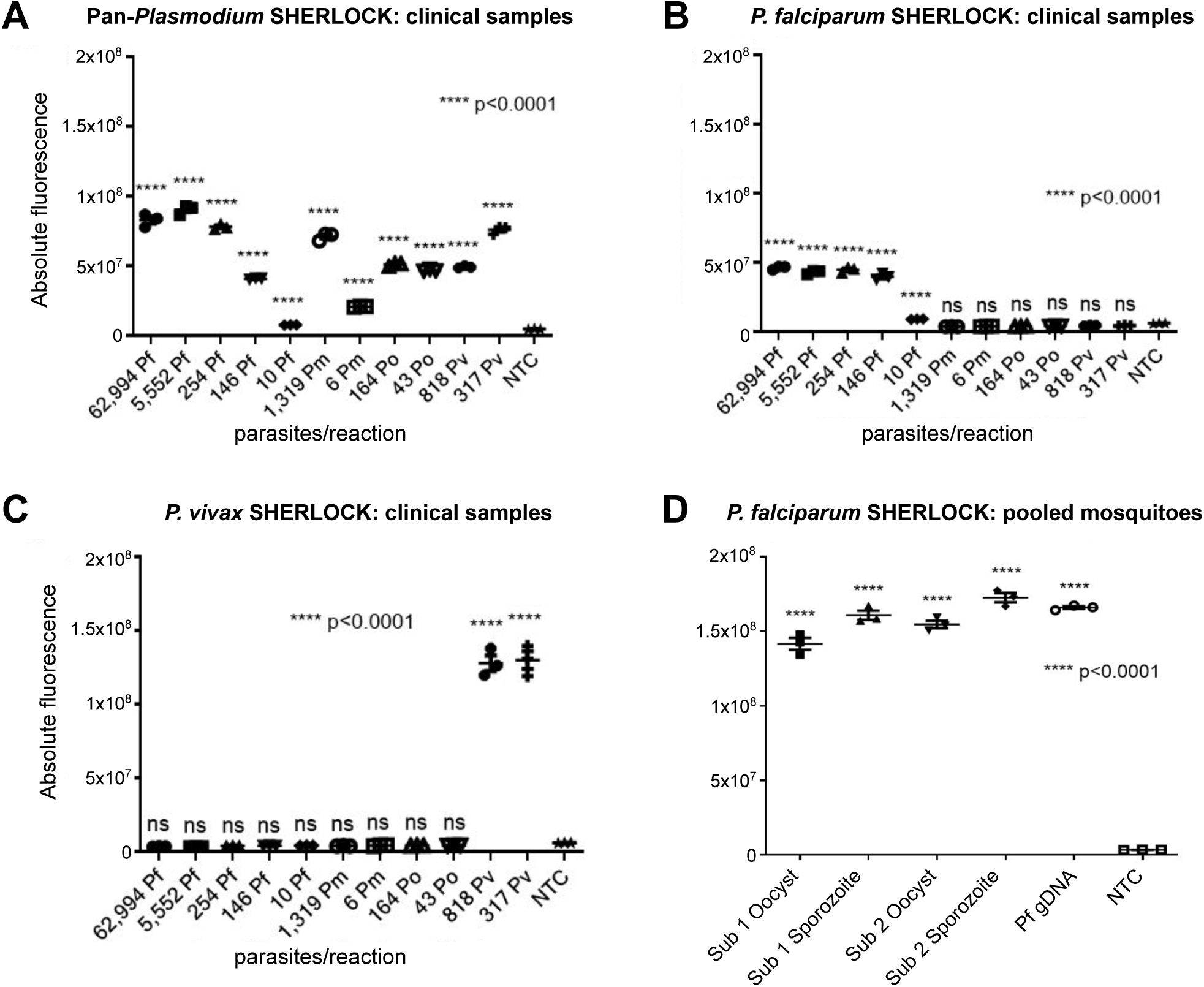
(**A**) *Plasmodium spp*., (**B**) *P. falciparum*, and (**C**) *P. vivax* SHERLOCK assays successfully detected and differentiated *Plasmodium* infections in diverse clinical isolates and (**D**) detected both the oocyst and sporozoite stages in infected mosquitos. Pf gDNA in the mosquito assay included 100,000 genome equivalents per reaction. Parasite densities were determined by qPCR for Pf, Pm, and Po and by microscopy for Pv. Abbreviations: Pf, *P. falciparum;* Pv, *P. vivax*; Pm, *P. malariae;* Po, *P. ovale*; NTC, no template control; gDNA, genomic DNA; Sub, subject.

**Figure 6.**
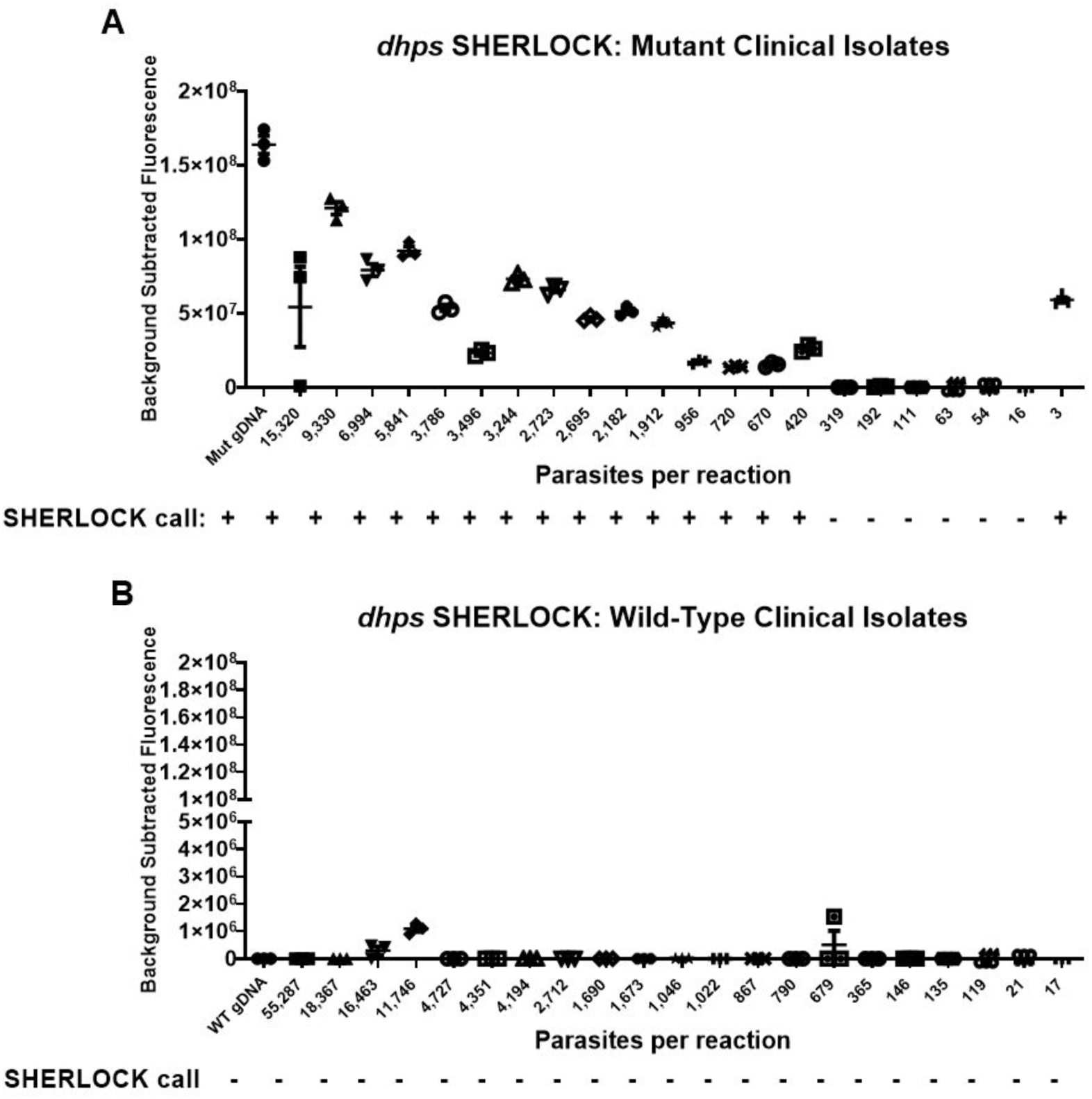
The *dhps* SHERLOCK assay detects *P. falciparum* parasites with (**A**) mutant (581G) but not (**B**) wild-type (A581) alleles. Parasite densities were determined by qPCR. Mutant (Mut) and WT gDNA standard inputs were 100,000 parasites/reaction.

The *P. falciparum* SHERLOCK assay also performed well when applied to DNA extracted from pooled, infected mosquitoes. In all cases, the *P. falciparum* SHERLOCK assay was able to detect the presence of parasites, including both the sporozoite and oocyst stages within the sporogonic life cycle.

### SHERLOCK can be used to detect SNVs associated with antimalarial drug resistance

Amplicon-based deep sequencing was performed to distinguish wild-type and A581G mutant *dhps P. falciparum* infections in 185 samples from the DRC and Uganda collected from patients with symptomatic malaria. The A581G mutation is associated with resistance to sulfadoxine-pyrimethamine, a drug commonly used to prevent malaria in pregnancy which can lead to poor birth outcomes. The median read count per sample was 81,775. After filtering, only two haplotypes were observed across all samples, corresponding to wild-type and A581G mutant *dhps*. Forty-three samples that were confirmed to be mono-infections bearing only wild-type or A581G mutant *dhps* mutations were selected for testing by SHERLOCK.

When applied to 22 wild-type and 21 A581G mutant *dhps* samples from Uganda, the sensitivity and specificity of the *P. falciparum dhps* SHERLOCK assay were 73% and 100%, respectively, using *dhps* deep sequencing calls as the gold standard. Cohen’s kappa was 0.72, consistent with good agreement between SHERLOCK and amplicon-based deep sequencing. When restricted to field samples with parasite densities of ≥420 parasites/µL, the *dhps* SHERLOCK assay’s clinical sensitivity and specificity were 100% and 100%, respectively. Among these higher parasite density samples, Cohen’s kappa was 1.0, consistent with perfect agreement between the *dhps* SHERLOCK assay and amplicon-based deep sequencing in this subset.

## DISCUSSION

By creating a new repertoire of SHERLOCK malaria assays, we highlight the versatility of CRISPR-based diagnostics and their potential to quickly improve the surveillance of important infectious diseases. They achieved robust clinical sensitivity and specificity when applied to well-characterized clinical samples and were easily adapted for parasite detection in mosquitoes and for drug-resistance genotyping. New diagnostics that can be used to detect, speciate, and genotype *P. falciparum* infections rapidly and reliably are especially needed in sub-Saharan Africa, where 94% of malaria deaths occur.*(12)*

SHERLOCK malaria assays outperform HRP2-based RDTs, the predominant malaria diagnostic modality deployed throughout Africa, and achieve similar performance to recently developed ultrasensitive HRP2-based RDTs, which have achieved clinical sensitivities of roughly 3 parasites/µL.*(13, 14)* Increasing reports of *P. falciparum* with deletion mutations of the histidine rich 2 and/or 3 (*pfhrp2/3*) genes raise concerns about reliance on HRP2 as the primary diagnostic target for RDTs.*(7)* RDTs that detect alternative antigens such as pLDH are available, but they are less sensitive and less heat-stable than HRP2-based RDTs. Thus, there is a need for novel malaria diagnostics that detect other targets. While both SHERLOCK and other CRISPR-based diagnostic assays currently in development need additional optimization prior to field deployment, we demonstrate that they are easily adapted for specific use cases and represent a promising avenue for malaria diagnostic development.

The analytical and clinical sensitivity of these SHERLOCK malaria assays approached that of commonly used real-time PCR assays, but the ability to conduct SHERLOCK without thermocycling allows for reduced laboratory infrastructure requirements and enables simplified isothermal approaches. More recently developed “ultrasensitive PCR” assays that target RNA and/or multicopy genes have achieved analytical sensitivities as low as 0.02p/µL, but their use in low-resource laboratory settings is limited by the need to protect against RNA degradation and/or the requirement for large-volume samples.*(15, 16)* Loop-mediated isothermal amplification (LAMP) is an alternative nucleic acid detection method that has been used for malaria diagnosis in field settings with similar sensitivity and specificity to SHERLOCK.*(17)* LAMP shows promise as a field-deployable molecular diagnostic, but its use for bespoke applications has been hampered by the complexity of target selection and primer design. CRISPR-based diagnostic approaches like SHERLOCK are an emerging technology that can be rapidly adapted in response to malaria’s evolving epidemiology while maintaining robust performance characteristics.*(1)*

Species discrimination was excellent across *Plasmodium spp*. SHERLOCK assays, with the exception of the *P. vivax* SHERLOCK assay that demonstrated low-level cross-reactivity with *P. knowlesi*. The *P. vivax* spacer and the analogous region on *P. knowlesi* differ by four nucleotides, so this cross-reactivity was unexpected. One explanation is that these nucleotides are towards the 3’ end of the crRNA spacer (positions 20, 26, 27 in the 28), which have been shown to have a smaller influence on crRNA binding efficiency.*(2)* Another explanation is unappreciated homology between *P. vivax* and *P. knowlesi* spacer binding sites in the setting of incomplete understanding of *P. knowlesi’*s genetic diversity. These findings highlight the importance of using high-quality genomic data when designing target sequences for CRISPR-based diagnostics and consideration of sequence variability within targets. Future SHERLOCK assays will be strengthened by efforts to improve our understanding of parasite genomic diversity.*(18, 19)*

To our knowledge, the novel SNV detection SHERLOCK assay for the *dhps* 581G variant in *P. falciparum* represents the first use of SHERLOCK to detect SNVs in clinical isolates outside of its original description.*(2, 3)* We chose to develop a SHERLOCK SNV detection assay for *Plasmodium dhps* variants due to their public health importance in the prevention of malaria during pregnancy. Variants in the *dhps* gene can confer resistance to sulfadoxine-pyrimethamine (SP), the primary antimalarial drug used in intermittent preventive treatment in pregnancy (IPTp) against placental malaria throughout much of sub-Saharan Africa.*(20)* Our *dhps* SHERLOCK assays demonstrated parasite-density dependent sensitivity and perfect specificity when compared to amplicon-based deep sequencing. When applied to a wide range of parasite inputs, the assay only produced false-negative results in samples with low parasite densities (6/7 samples with ≤319 parasites/µL sample input). While the SNV detection assay is concentration dependent, it performed well at parasite densities typically associated with clinically significant malaria.*(21)*

In future, mixed infections involving both wild-type and 581G mutant strains could be identified by pairing the 581G-specific assay described here with a wild-type-specific assay in a multiplexed format. SHERLOCK’s SNV detection capabilities are a promising tool for malaria control programs in low-resource settings where sequencing facilities are not available to support surveillance. For example, high prevalence of resistance-associated *dhps* variants detected by SHERLOCK could trigger deployment of alternative drug regimens for IPTp in affected regions.

SHERLOCK is now one of several CRISPR-based diagnostic methodologies; the Cas12a-based DETECTR operates similarly but instead detects dsDNA targets and cleaves single-stranded DNA (ssDNA) during collateral activation. We chose SHERLOCK over DETECTR for our study in large part because of Cas13a’s minimal protospacer-adjacent motif (PAM) site nucleotide requirements compared to Cas12a. LwCas13a requires only H (not G) adjacent to the spacer region, which makes guide design and SNV detection for SHERLOCK easier than for DETECTR, which has a PAM requirement of “TTTN”.*(22, 23)* Additionally, multiplexing with SHERLOCK has been demonstrated using Cas13a isolated from multiple bacterial species, which have different collateral activities that can be used to activate different RNA reporters, enabling multiplexed assays.*(3)* These properties provide opportunities for further assay development, including translation of our malaria SHERLOCK assays into a single, multiplexed assay that enables speciation of multiple *Plasmodium* species and drug-resistance SNVs in a single reaction.

We experienced success using SHERLOCK to detect *Plasmodium spp*. and drug-resistance variants; however, several limitations must be overcome for successful translation of SHERLOCK from the laboratory to the field. First, we could not achieve the sub-attomolar limits of detection or “one-pot” reaction conditions previously described for viral targets.*(3)* We also observed that our assays performed best using higher crRNA concentrations than those originally described.*(2)* Though this may reflect differences in crRNA synthesis or lot-to-lot variation in other reagents, we observed improved reaction performance with higher crRNA concentrations across multiple experiments. These obstacles did not prevent us from adapting this method for malaria, but they suggest that implementation of SHERLOCK is not yet “plug-and-play” but requires a degree of specialized experience. Second, the cost of designing and validating a SHERLOCK assay is not trivial. In a reaction volume of 25 µL used for 96-well plates and using fluorescent output, the cost was roughly equivalent to PCR at over $2.00 of reagent costs per technical replicate, plus upfront costs of crRNA optimization and synthesis and the need for a fluorescence plate-reader. These costs can be reduced for assays brought to scale but are an important consideration during assay design. Finally, SHERLOCK requires multiple reagents, custom crRNAs, and LwCas13a. Recent progress has been made to reduce its complexity and make these assays more easily accessible.*(24)* Implementation of SHERLOCK as a point-of-care molecular diagnostic will require further streamlining, including the development of commercially-available mastermixes and the combination of nucleic acid extraction, LwCas13a activation, and signal readout into a single device. Recent reports suggest that these advances are within reach.*(4, 25)*

While these limitations must be overcome before immediate field applications, SHERLOCK remains a promising technology for sensitive and specific pathogen detection. Lateral flow platforms and miniaturization promise to lower the cost per reaction. SHERLOCK’s use for SNV detection and multiplexing are possible niches for use in a variety of settings, including both surveillance and diagnosis of infectious diseases. Continued development of new Cas effectors, streamlined workflows, and point-of-care read-outs will open new opportunities for CRISPR diagnostics in clinical, surveillance, and research applications. These novel malaria SHERLOCK assays confirm the promise of CRISPR-based diagnostics for diverse applications and in resource-poor settings.

## MATERIALS AND METHODS

### RPA primer design

We designed RPA primers capable of genus-specific amplification of the *Plasmodium* 18S ribosomal RNA gene, building upon existing TaqMan real-time PCR assays.*(10)* PCR primers were extended to 30-35 nt in length and modified to include nucleotide favorability criteria described in the TwistAmp Assay Design Manual.*(26)* These RPA primers target regions of the 18S rRNA gene that are conserved across human-infecting *Plasmodium* species, allowing “plug and play” crRNA design for speciation during the crRNA:LwCas13a base-pairing step of SHERLOCK. RPA primers for the *dhps* SNV-detection SHERLOCK assays were designed using NCBI Primer-BLAST with default parameters and the following modifications: amplicon size 100-140 nt, melting temperatures between 54-67°C, and length between 30 and 35 nucleotides.*(27)* We then added a T7 promoter sequence (5’-GAAATTAATACGACTCACTATAGGG-3’) to the 5’ end of one RPA primer in each set to enable *in vitro* transcription by T7 polymerase during the LwCas13a detection step. All primers with off-target complementarity predicted by Basic Local Alignment Search Tool (BLAST) were excluded. Salt-free primers were synthesized by Integrated DNA Technologies (Coralville, Indiana). The full list of oligonucleotides designed and used in this manuscript can be found in **Supplementary Table 1**.

### crRNA design

Oligonucleotides for crRNA synthesis were designed as two complementary ssDNA oligonucleotides and then *in vitro* transcribed to produce single-stranded 67 nt crRNAs. Each ssDNA oligonucleotide was composed of three parts: a variable spacer region (to facilitate recognition of the RNA target molecule), a constant region (to facilitate crRNA association with LwCas13a), and a T7 promoter sequence (to facilitate crRNA *in vitro* transcription).

Spacers are 28 nt long and recognize the *in vitro* transcribed RNA product of the RPA reaction through complementary base-pairing.*(2)* For the species-specific *Plasmodium* SHERLOCK assays, spacers were designed to bind a variable but species-specific region of the 18S rRNA gene previously targeted for amplification during real-time PCR assays.*(10)* Spacers were selected to have at least three mismatches between species for the *P. falciparum* and *P. vivax*-specific assays to minimize off-target activity. For the pan-*Plasmodium* SHERLOCK assay, a spacer was selected in a region of the 18S rRNA gene conserved between *Plasmodium* species.

Spacers for the *dhps* SHERLOCK assays were designed to maximize the difference in signal between a specific SNV and alternate motifs as previously described.*(2)* To accomplish this, spacers were designed so the SNV is recognized at position 3 or 6, and a synthetic mismatch between the spacer and target was present at position 5 or 4, respectively **(Supplementary Table 2)**. The resulting spacer has one mismatch with the SNV of interest and two mismatches with the alternate variant, producing a measurable difference in signal between the two variants **(Figure 2)**. Spacers were queried through Basic Local Alignment Search Tool (BLAST) to search for any off-target complementarity.*(27)*

The same constant LwCas13a-associating region was used as described in Gootenberg et al. (2017),*(2)* 5’-GGGGAUUUAGACUACCCCAAAAACGAAGGGGACUAAAAC-3’, and the T7 promoter sequence used was 5’-GAAATTAATACGACTCACTATAGGG-3’. Put together, two 89 nt ssDNA oligonucleotides (appended to spacer sequences) for each crRNA were designed in the format 5’-GAAATTAATACGACTCACTATAGGGGATTTAGACTACCCCAAAAACGAAGGGGACT AAAAC-spacer-3’ and 5’-spacer reverse complement-GTTTTAGTCCCCTTCGTTTTTGGGGTAGTCTAAATCCCCTATAGTGAGTCGTATTAA TTTC-3’. These oligonucleotides were ordered from Integrated DNA Technologies (Coralville, IA). For each SHERLOCK assay, multiple crRNA designs were screened, and crRNAs with the greatest signal:background and/or SNV signal:alternate signal were selected for all future experiments **(Supplemental Figures 1-3)**.

### crRNA synthesis

Complementary ssDNA oligonucleotides were annealed (final concentration 10 µM each) in 10 mM Tris-HCl, 50 mM KCl, and 1.5 mM MgCl_2_ pH 8.3 for 5 minutes at 95°C followed by a 5°C/minute temperature decrease to 25°C. dsDNA templates were then *in vitro* transcribed using the HiScribe T7 Quick High Yield RNA Synthesis Kit (New England Biolabs, Ipswich, MA) using overnight incubation as described in the protocol for <0.3kb transcripts.*(28)* CrRNAs were purified using RNAClean XP magnetic bead purification (Beckman Coulter, Indianapolis, IN) and quantified by NanoDrop spectrophotometry (ThermoFisher, Waltham, MA).

### LwCas13a synthesis and purification

LwCas13a protein was synthesized and purified at the University of North Carolina at Chapel Hill (UNC) Protein Expression and Purification Core as previously described with several modifications.*(2)* Briefly, the pET His6-TwinStrep-SUMO-LwCas13a (NovoPro, Shanghai, China) expression vector was transformed into Rosetta™ 2 (DE3) pLysS Singles Competent Competent Cells (Millipore) in autoinduction media and grown at 37°C until OD=0.6, when the temperature was lowered to 18C and grown overnight. The pellet was harvested and stored at -80°C until purification. For purification, the pellet was resuspended in lysis buffer (50 mM NaPO_4_, 500 mM NaCl, 20 mM Imidazole, 1 mM PMSF, 50 µg/mL lysozyme) and mixed at 4°C for 30 min. The solution was then sonicated on ice and centrifuged for 1 hour at 10,000 x g at 4°C. Clarified supernatant was filtered using a 0.45 micron filter. Filtered supernatant was then loaded over Nickel Sepharose 6 Fast Flow resin (GE). The column was washed with 5 CV Buffer A (50 mM NaPO_4_, 500 mM NaCl, 20 mM imidazole) and eluted with 10 CV Buffer B (50 mM NaPO_4_, 500 mM NaCl, 500 mM imidazole). Buffer exchange was performed via dialysis to a final solution of 600 mM NaCl, 50 mM Tris-HCl pH 7.5, 5% glycerol, and 2mM DTT.

### Recombinase polymerase amplification

RPA reactions were conducted as described in TwistAmp Basic Instruction Manual (TwistDx, Cambridge, UK) with several modifications.*(29)* Each 50 µL TwistAmp reaction was subdivided by first creating a 45 µL mastermix and aliquoting five 5 × 9 µL reactions before adding 1 µL of sample input. Magnesium acetate was added to the mastermix before adding the sample input. To simulate human background in clinical samples, 40 ng human gDNA from buffy coat (Sigma Aldrich, St. Louis, MO) was added in all experiments except those using clinical isolates. RPA reactions were incubated at 37 °C for 30 minutes in a thermal cycler with a 40 °C heated lid. After the first 4 minutes of incubation, samples were removed, vortexed briefly, and then returned to the thermal cycler for the remainder of the incubation period.

### Detection using LwCas13a

LwCas13a detection reactions were performed as previously described with several modifications. Each reaction was performed using 25 μL total volume and contained 45 nM LwCas13, 100-500 ng crRNA, 125 nM fluorescent RNA reporter (RNAse Alert v2, ThermoFisher), 0.625 μL murine RNase inhibitor (New England Biolabs), 31.25 ng background RNA (isolated from Burkitt’s Lymphoma (Raji), ThermoFisher), 1 mM rNTP mix (New England Biolabs), 0.75 μL T7 polymerase (New England Biolabs), 3mM MgCl_2_, and 1.25 uL of RPA product in nuclease assay buffer (20 mM HEPES, 60 mM NaCl, 6 mM MgCl2, pH 6.8).*(3)* Detection reactions were conducted in 96-well black half-area microplates (PerkinElmer, Waltham, MA) and sealed with MicroAmp optical adhesive film (ThermoFisher). Detection reactions were incubated for 3 hours at 37°C on a VICTOR Nivo fluorescent plate reader (PerkinElmer) with fluorescence measurements taken every 5 minutes. Background subtracted fluorescence for each sample was calculated by subtracting the average fluorescence intensity of the human gDNA controls at 180 minutes of the human gDNA control from each sample.

### Analytical sensitivity and specificity estimation

To determine the analytical sensitivity of the SHERLOCK assays, we performed multiple replicates of each assay, using serially diluted template DNA. For these experiments, we used serially diluted genomic DNA from *P. falciparum* strain Dd2 (MRA-150G) for *P. falciparum, P. vivax* 18S rRNA plasmid DNA (MRA-178), *P. ovale* 18S rRNA plasmid DNA (MRA-180), P. malariae 18S rRNA plasmid DNA (MRA-179), and P. knowlesi genomic DNA (MRA-456G) (Bei Resources, Manassas, VA). To facilitate ease of interpretation, we assumed 6 copies of 18S rRNA targets per genome when calculating parasite genome equivalents.*(30)*

We first tested the analytical performance of assays in triplicate using serially diluted parasites ranging from 10^5^ and 10^−2^ parasite genomes/µL. This 10-fold dilution series was selected to include expected parasite densities observed during human infection, including clinical malaria (typically 10^2^ to 10^5^ parasites/µL) and subclinical infection. Differences in mean background-subtracted fluorescence were assessed using the student’s unpaired t-test in Prism (GraphPad, San Diego, CA). We then performed additional biological replicates using 2-fold serial dilutions near the observed limit of detection to improve the precision of our analytical sensitivity estimates. We used probit analysis to determine the 95% limit of detection and 95% confidence intervals for each SHERLOCK assay. Data analysis was conducted in R (R Core Team, Vienna, Austria). Finally, we assessed each assay’s analytical specificity using high-concentration DNA for all human-infecting *Plasmodium* species, including gDNA and 18S rRNA plasmid DNA as described above and containing 100,000 genome equivalents per reaction.

### Clinical sensitivity and specificity estimation

We assessed the clinical sensitivity and specificity of our assays using a panel of dried blood spot (DBS) samples collected in the DRC, Uganda, and Thailand. A summary of the sources of clinical samples is provided in **Supplementary Table 3**. DRC samples were collected from subjects presenting to government health facilities with symptoms of malaria in Kinshasa, South Kivu, and Bas-Uele Provinces in 2017 as part of a separate study of malaria rapid diagnostic test performance.*(31)* Ugandan samples were collected from febrile children presenting to public health facilities located in the Kasese District of western Uganda over the period November 2017 to June 2018.*(32)* Thai samples were collected from patients presenting to public health clinics and found to be smear-positive for *P. vivax* and as part of ongoing entomological surveillance in the Mae Sod district of Tak Province, northwest Thailand in 2010. Ethical approvals were obtained by the Kinshasa School of Public Health, Mbarara University of Science and Technology, Uganda National Council for Science and Technology, Walter Reed Army Institute of Research, Ministry of Public Health in Thailand, and the University of North Carolina at Chapel Hill. All enrolled subjects provided informed consent.

All samples were speciated using real-time PCR. In brief, DNA was extracted from three 6mm DBS punches using Chelex-100 and saponin for DRC samples,*(33)* Chelex-100 and Tween for Uganda samples,*(34)* and the QIAamp DNA mini kit (QIAGEN, Hilden, Germany) for Thailand samples prior to storage at -20°C. DRC samples were first subjected to a pan-*Plasmodium* real-time PCR assay targeting the 18S rRNA gene.*(35)* Positive samples were then subjected to four species-specific, semi-quantitative 18S rRNA real-time PCR assays for *P. falciparum,(36) P. ovale,(37) P. malariae,(36)* and *P. vivax,(38)* respectively. Parasite densities for all DRC and Uganda samples were determined using quantitative real-time PCR (qPCR) targeting the single-copy *P. falciparum* lactate dehydrogenase (*pfldh*) gene as previously described.*(39, 40) P. vivax* infection was diagnosed and parasite densities determined in Thailand using microscopy at the time of DBS collection and later confirmed using qualitative, species-specific 18S rRNA real-time PCR for *P. vivax*.*(36, 38)* PCR primers, probes, and reaction conditions are described in the **Supplementary File**.

We first assessed the performance of all three assays using a panel of 11 DNA samples from real-time PCR-confirmed *Plasmodium* infections, including all species and a range of parasite densities. We further characterized the performance of the *P. falciparum* assay using 112 samples collected in the DRC and Uganda. All *Plasmodium* SHERLOCK assays were performed in triplicate. Clinical sensitivity, clinical specificity, and Cohen’s Kappa coefficient were calculated using real-time PCR as the gold standard.

### Parasite DNA detection in mosquito pools using SHERLOCK

Parasite DNA was extracted from insectary-reared *Anopheles dirus* mosquitoes fed on blood from gametocytemic *P. falciparum*-infected volunteers in Cambodia. Mosquitoes were saved 9 and 16 days post-feeding to capture oocyst-stage and sporozoite-stage infection, respectively. They were pooled in groups of 10 and preserved in 95% ethanol before undergoing DNA extraction via a simplified chelex protocol as previously described.*(41)* SHERLOCK assays were performed on DNA from mosquito pools in triplicate. Comparison of mean background-subtracted fluorescence values was performed using the t-test as described above.

### Deep sequencing of clinical samples and dhps genotyping

We selected 463 DNA samples from the DRC with *P. falciparum* mono-infection by real-time PCR for *dhps* genotyping using a multiplex real-time PCR assay that discriminates wild-type from 581G mutants (see **Supplementary File** for details).*(42)* We then selected samples for amplicon-based deep sequencing, including a subset of 92 wild-type and 581G mutant DNA samples identified during *dhps* real-time PCR screening of DRC samples and 92 *P. falciparum* histidine-rich protein 2-based RDT-positive samples from Uganda. Each of these 184 samples was amplified using a barcode-labeled forward and reverse primer (**Supplementary Table 4**). PCR reactions were carried out in a total volume of 50µL and using 5µL of input DNA. The reaction mixture contained 1X Accuprime PCR buffer II (Invitrogen, Carlsbad, CA), 1 unit Accuprime HiFi taq, 1µL of 0.0025 mg/µL BSA, and 400nM of forward and reverse primer. The reaction conditions were 95°C for 2 minutes, followed by 40 cycles of 95°C for 30 seconds, 60°C for 30 seconds and 68°C for 1 minute, with a final extension of 68°C for 10 minutes. PCR products were quantified using picogreen on a microplate reader. Amplicons were pooled, prepared for sequencing using Kappa HyperPrep Kits (Roche, Indianapolis, IN) and sequenced on a single MiSeq (Illumina, San Diego, CA) run using 250bp paired-end chemistry at the UNC High Throughput Sequencing Facility.

Sequencing data was analyzed using SeekDeep v2.6.5 using default parameters.*(43)* In brief, data was demultiplexed using the *pfdhps* primers and a dual barcoding scheme, and final haplotypes for each sample were determined by removing low per-base quality scores and low frequency mismatches. Final haplotypes were further filtered by removing haplotypes with <2,000 reads within samples. We then selected DNA from samples containing a single haplotype consistent with a wild-type or A581G mutant parasite for SHERLOCK SNV testing. Mixed infections were excluded. Wild-type parasites were stratified by sampling location and selected for SHERLOCK testing using a random number generator (Excel, Microsoft, Seattle, WA).

### Dhps SNV detection by SHERLOCK

SHERLOCK can detect SNVs by differential crRNA binding and LwCas13a activation. Briefly, crRNA-target RNA binding interactions can typically tolerate a single base-pair mismatch in the early spacer region, resulting in high-moderate fluorescence output **(Figure 2)**. However, a second mismatch greatly decreases crRNA binding and LwCas13a activation, resulting in significantly lower fluorescence output. By introducing a synthetic mismatch in the crRNA spacer design, the presence or absence of the SNV will cause large changes in fluorescent signal. All *dhps* SNV-detection SHERLOCK assays were performed in triplicate. Positive SHERLOCK calls were made if the median absolute fluorescence was at least 20% above the mean background fluorescence. Clinical sensitivity, clinical specificity, and Cohen’s Kappa coefficient were calculated using deep sequencing calls as the gold standard.

## SUPPLEMENTARY MATERIALS

**Supplementary Figure 1:**
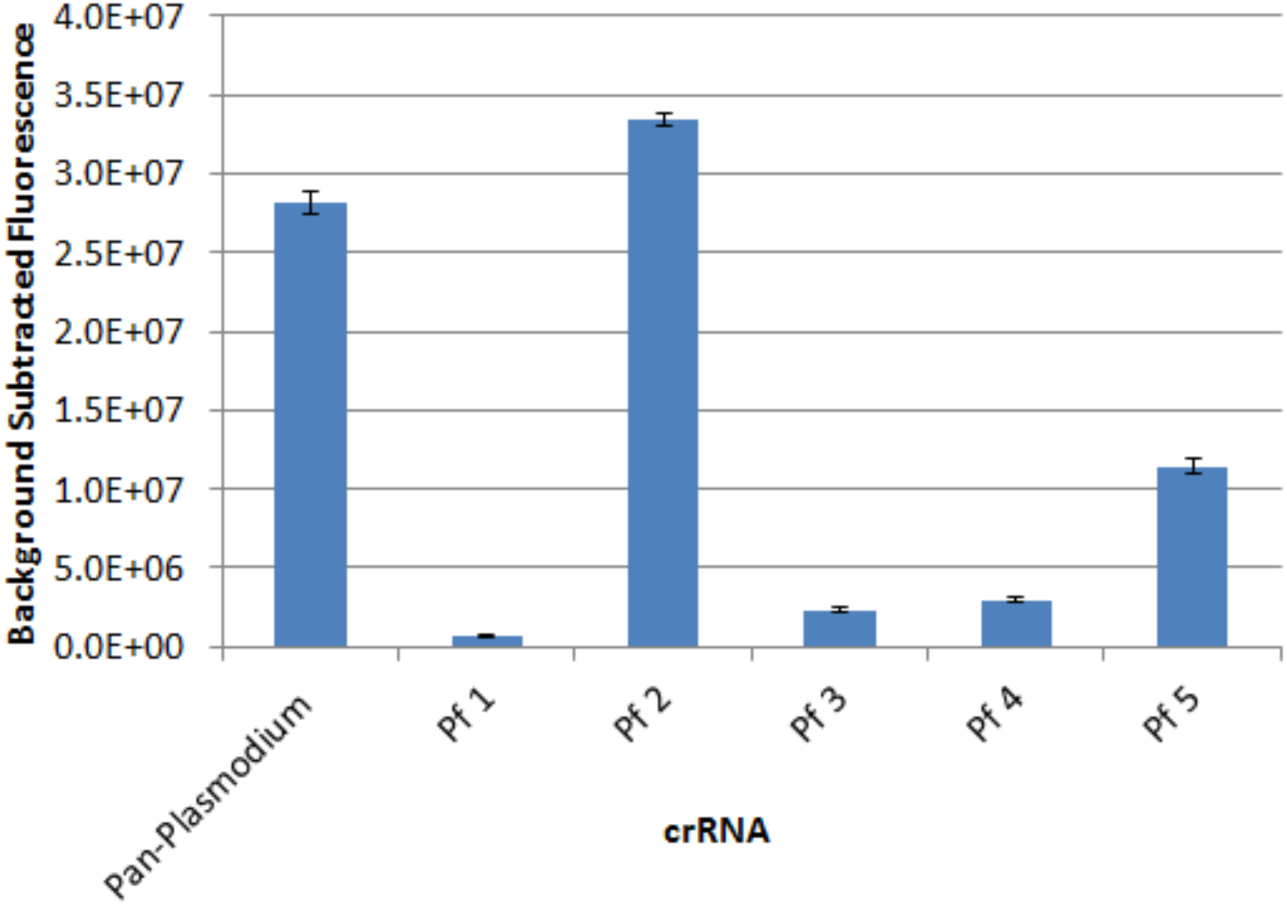
*P. falciparum* crRNA screen. Pf 2 was selected for further testing based on its background-subtracted fluorescence. Input: *P. falciparum* 18S rRNA plasmid, 100,000 genome-equivalents/reaction. Bars depict mean results ± standard error.

**Supplementary Figure 2:**
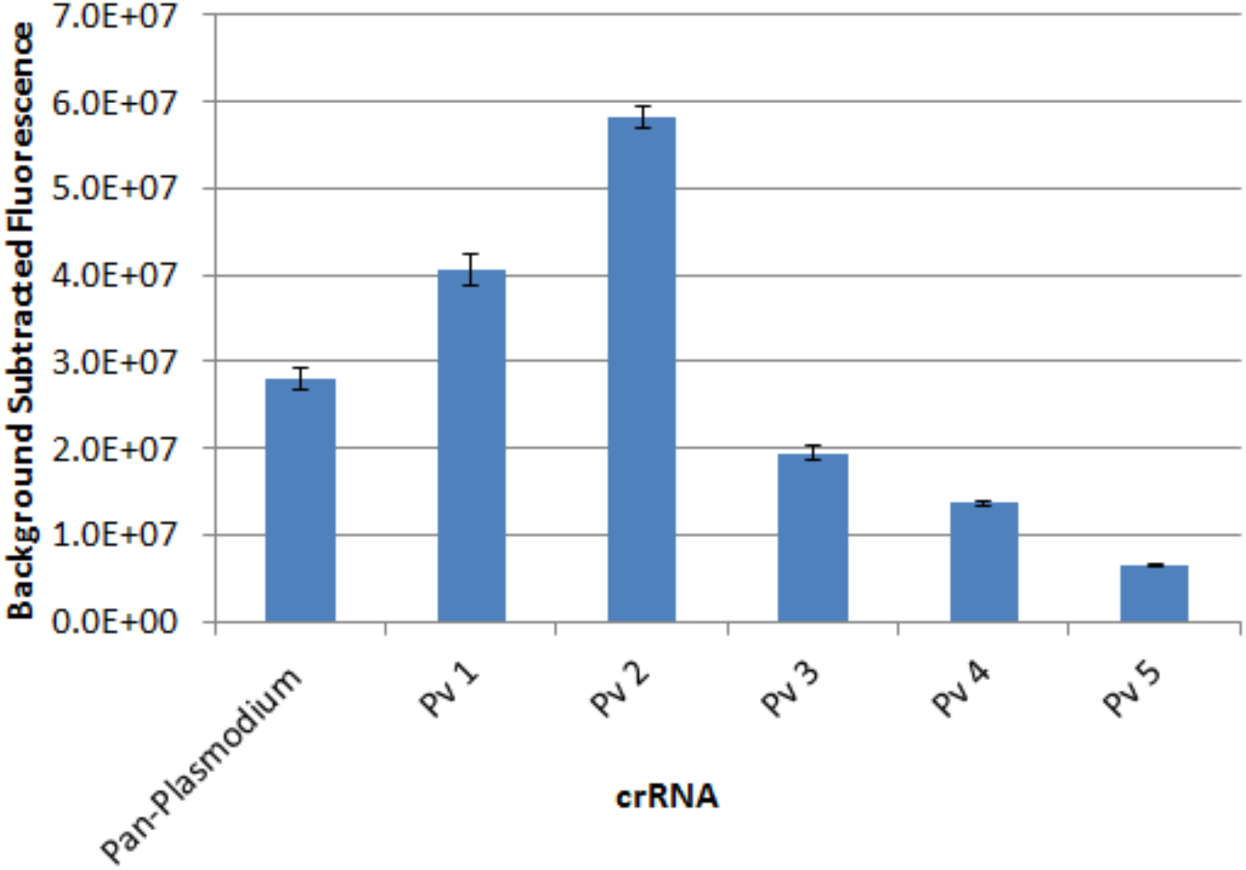
*P. vivax* crRNA screen. Pv 2 was selected for further testing based on its background-subtracted fluorescence. Input: *P. vivax* 18S rRNA plasmid, 100,000 genome-equivalents/reaction. Bars depict mean results ± standard error.

**Supplementary Figure 3:**
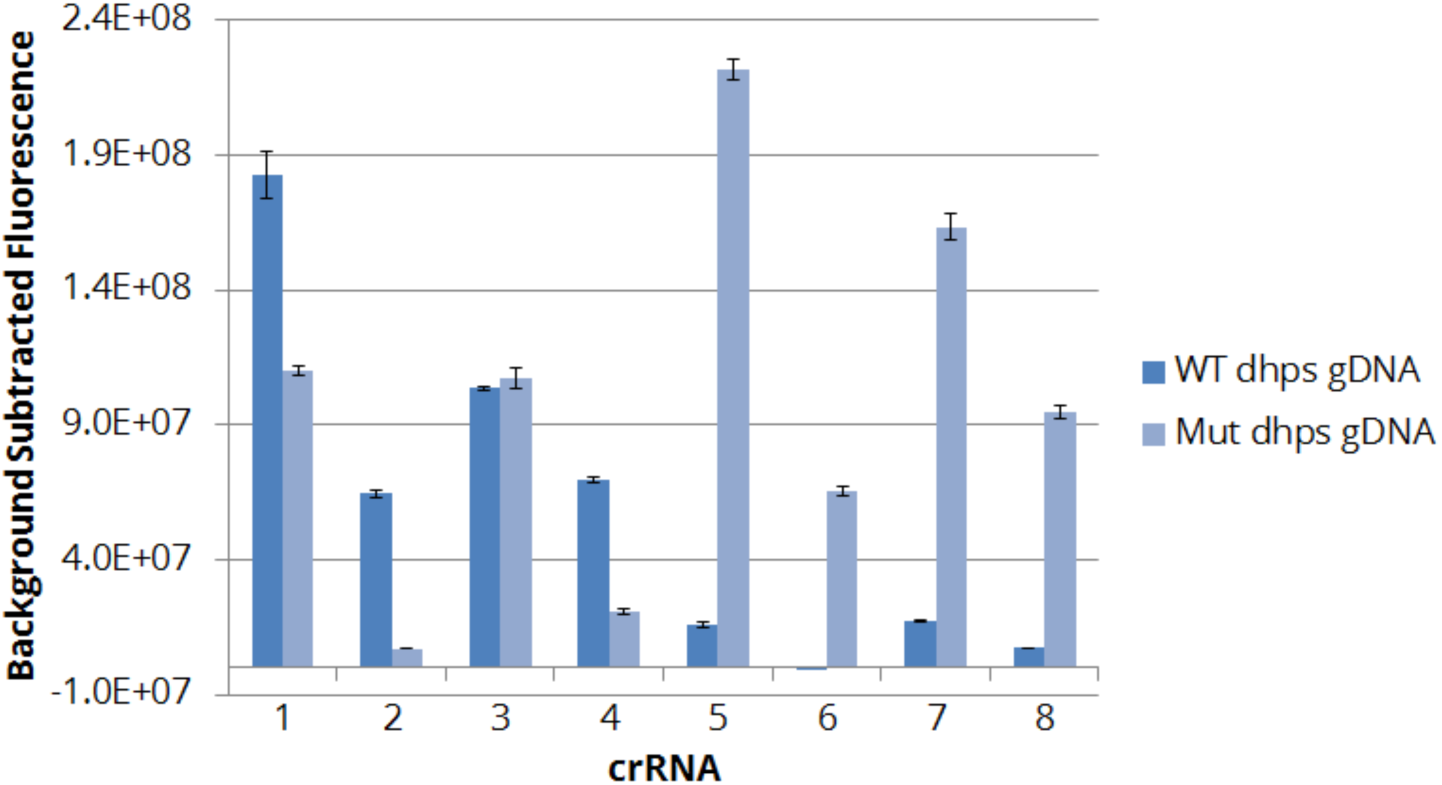
Dhps crRNA screen. *Dhps* crRNA 5 was selected for further testing in the 581G SNV-detecting SHERLOCK assay based on its high mutant:wild-type background-subtracted fluorescence ratio. Wild-type (WT) gDNA: Dd2 strain *P. falciparum*, 100,000 copies/reaction. Mutant (Mut) gDNA: K1 strain *P. falciparum*, 100,000 copies/reaction. Bars depict mean results ± standard error.

**Supplementary Figure 4:**
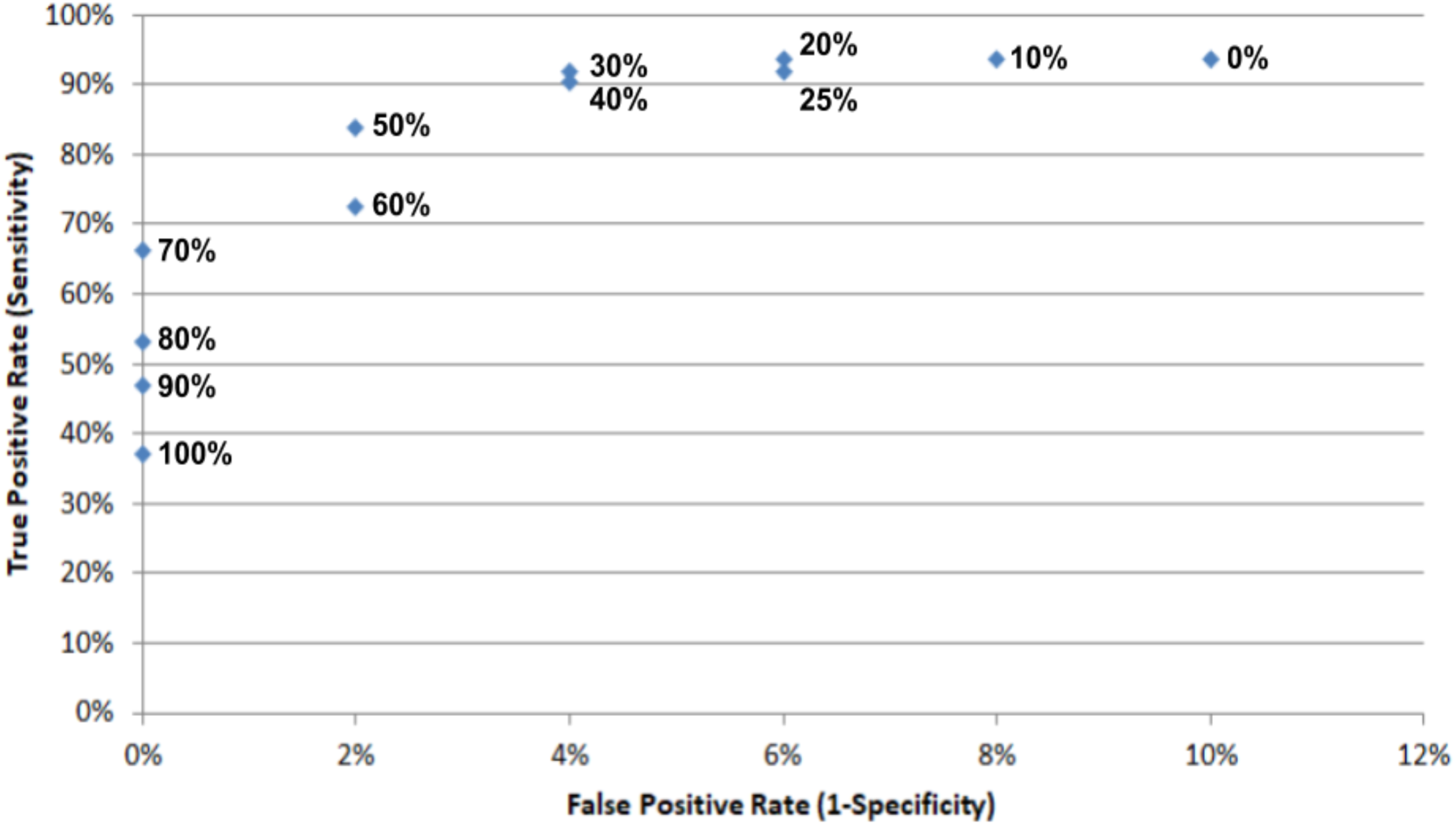
Receiver-operator curve for *P. falciparum* SHERLOCK assay tested on 62 real-time PCR-positive and 50 real-time PCR-negative clinical isolates. Points represent candidate thresholds for a positive SHERLOCK call, using the formula *(absolute fluorescence of sample)/(absolute fluorescence of negative control)x100* to calculate threshold percentages. 20% signal above background was selected as the cutoff for a positive call, yielding 94% sensitivity and 94% specificity compared to real-time PCR as the gold standard.

**Supplementary Table 1:**
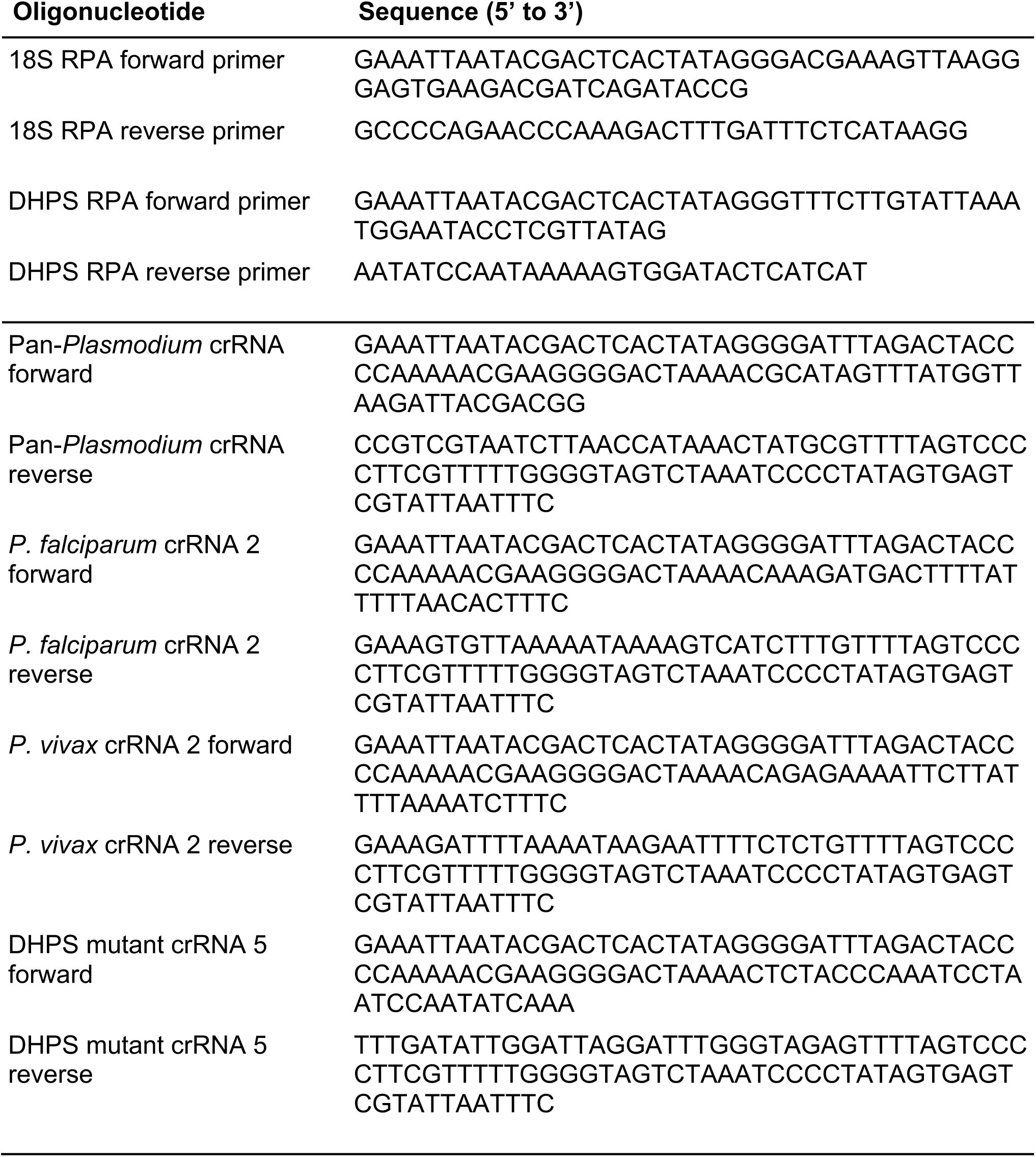
Oligonucleotides used as RPA primers and DNA templates for crRNA synthesis.

**Supplementary Table 2:** crRNAs used in *dhps* crRNA screen.

| Detector type | crRNA | WT mismatches | Mut mismatches | Mismatch positions |
| --- | --- | --- | --- | --- |
| Wild-type | 1 | 0 | 1 | 3,5 |
|  | 2 | 1 | 2 | 3,5 |
|  | 3 | 0 | 1 | 4,6 |
|  | 4 | 1 | 2 | 4,6 |
| Mutant | 5 | 1 | 0 | 3,5 |
|  | 6 | 2 | 1 | 3,5 |
|  | 7 | 1 | 0 | 4,6 |
|  | 8 | 2 | 1 | 4,6 |

**Supplementary Table 3:** Sources of clinical isolates. Abbreviations: DRC, Democratic Republic of the Congo.

| Experiments | Clinical isolates | Number | Source |
| --- | --- | --- | --- |
| Clinical sample speciation<br>*Fig. 5A-C* | *P. falciparum* | 5 | DRC |
|  | *P. malariae* | 2 | DRC |
|  | *P. ovale* | 2 | DRC |
|  | *P. vivax* | 2 | Thailand |
| Mosquito DNA detection<br>*Fig. 5D* | Mosquito DNA extracts | 4 | Thailand |
| *dhps* SNV detection<br>*Fig. 6* | A581 *P. falciparum* | 22 | Uganda |
|  | 581G *P. falciparum* | 21 | Uganda |
| *P. falciparum* detection<br>*Supp. Fig. 4* | *P. falciparum*-positive | 62 | Uganda |
|  | *P. falciparum*-negative | 50 | DRC |

**Supplementary Table 4.**
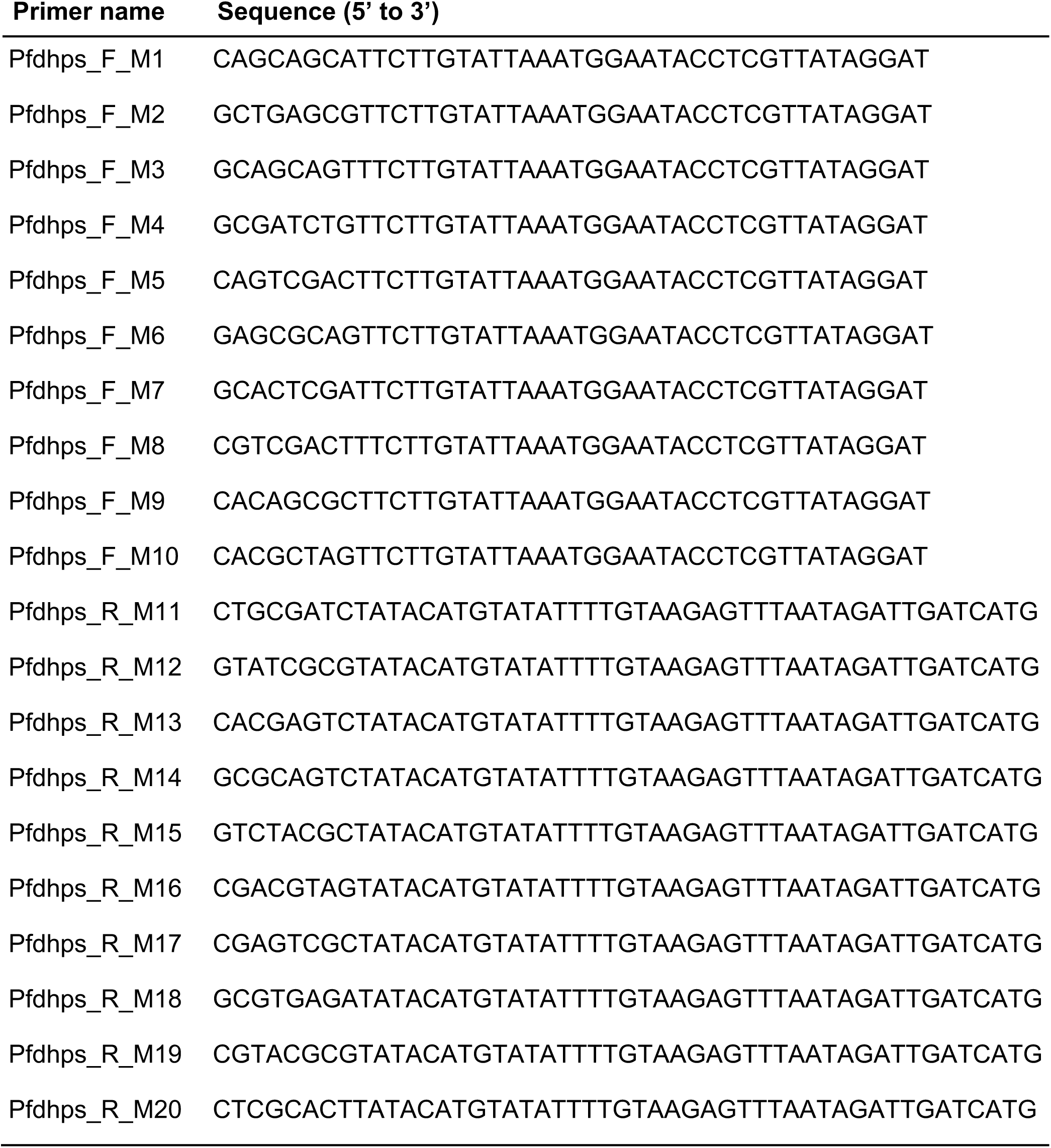
Primers used for *dhps* deep sequencing.

## Supplementary File

PCR primers, probes, and conditions used on samples in the study.

## FOOTNOTES

## Acknowledgements

The authors would like to thank the study coordinators and teams from SANRU Asbl and Kinshasa School of Public Health in the DRC, the Mbarara University of Science and Technology in Uganda, and the Armed Forces Research Institute of Medical Sciences in Thailand who conducted field work for the parent studies from which clinical samples and mosquito pools were selected. They also thank Dr. Qi Zhang for helping troubleshoot crRNA synthesis and storage; the UNC Protein Expression and Purification core for LwCas13a synthesis; Dr. Jeff Laux for biostatistical support; Drs. Jonathan Gootenberg, Omar Abudayyeh, and Feng Zheng for helpful advice during assay troubleshooting. Most importantly, the authors would like to thank the patients who provided samples that enabled assay validation.

The following reagents were obtained through BEI Resources, NIAID, NIH: Genomic DNA from *Plasmodium falciparum*, Strain Dd2, MRA-150G, contributed by David Walliker; Diagnostic plasmids containing the small subunit ribosomal RNA gene (18S) from *Plasmodium vivax*, MRA-178, *Plasmodium ovale*, MRA-180, and *Plasmodium malariae*, MRA-179, contributed by Peter A. Zimmerman; and Genomic DNA from *Plasmodium knowlesi*, Strain H, MRA-456G, contributed by Alan W. Thomas.

## Author contributions

CHC: Conceptualization, Methodology, Validation, Formal Analysis, Investigation, Visualization, Writing - Original Draft. CH, KLT: Investigation. JTL, RU, RMB, EMM, FP, AK, KM, AT, SRM: Resources. NH: Software, Formal Analysis. JJJ: Conceptualization, Supervision, Funding Acquisition. JBP: Conceptualization, Supervision, Project Administration, Funding Acquisition, Visualization, Resources, Writing - Original Draft. All authors: Writing - Review and Editing.

## Data and materials availability

All data associated with this study are available in the main text, the supplementary materials, or upon reasonable request to the authors.

## List of Supplementary Materials

1. Supplementary Materials: Figures and Tables
2. Supplementary File: PCR primers and conditions

